# Interspecific allometric scaling in eDNA production in fishes reflects physiological and surface area allometry

**DOI:** 10.1101/2022.04.22.489177

**Authors:** M.C Yates, T. W. Wilcox, M.Y. Stoeckle, D.D. Heath

**Affiliations:** University of Windsor, Windsor, Ontario, CA; National Genomics Center for Wildlife and Fish Conservation, Rocky Mountain Research Station, Missoula, MT, USA; The Rockefeller University, New York, NY, USA

**Keywords:** eDNA, abundance, allometry, allometric scaling, fishes, biomass

## Abstract

Relating environmental DNA (eDNA) signal strength to organism abundance requires a fundamental understanding of eDNA production. A number of studies have demonstrated that eDNA production may scale allometrically – that is, larger organisms tend to exhibit lower mass-specific eDNA production rates, likely due to allometric scaling in key processes related to eDNA production (e.g. surface area, excretion/egestion). While most previous studies have examined intra-specific allometry, physiological rates and organism surface area also scale allometrically across species. We therefore hypothesize that eDNA production will similarly exhibit inter-specific allometric scaling. To evaluate this hypothesis, we reanalyzed previously published eDNA data from Stoeckle et al. (2021) which compared metabarcoding read count to organism count and biomass data obtained from trawl surveys. Using a Bayesian model we empirically estimated the value of the allometric scaling coefficient (‘b’) for bony fishes to be 0.67 (credible interval = 0.58 – 0.77), although our model failed to converge for chondrichthyan species. We found that integrating allometry significantly improved correlations between organism abundance and metabarcoding read count relative to traditional metrics of abundance (density and biomass) for bony fishes. Although substantial unexplained variation remains in the relationship between read count and organism abundance, our study provides evidence that eDNA production tends to scale allometrically across species. Future studies investigating the relationship between eDNA signal strength and metrics of fish abundance could potentially be improved by accounting for allometry – a scaling coefficient value of ∼2/3 appears to be both theoretically and empirically justified.

## Introduction

A consensus is emerging that the relationship between the amount of DNA in an environment (environmental DNA, or ‘eDNA’) tends to be positively correlated with the abundance of organisms within that environment (Yates et al. 2019, 2021b; Rourke et al. 2021). The concentration of eDNA can therefore provide information on the ‘unseen’ abundance of organisms, and could represent a powerful tool for understanding aquatic community composition. However, studies in natural ecosystems exhibit substantial variation in the strength of the correlation between eDNA concentration and estimated organism abundance, likely because a number of abiotic and biotic variables can affect steady-state concentrations of eDNA (Yates et al. 2021b). Improving the capacity to infer abundance from eDNA data therefore requires a better understanding of the ‘ecology’ of eDNA – that is, factors influencing the production, transport, and degradation of nucleic acids in the environment (Barnes and Turner 2016; Yates et al. 2021b).

A number of previous studies have demonstrated that eDNA production likely scales allometrically with body size; that is, as the body mass of an individual increases, their *mass-specific* eDNA production rate (*i*.*e*. production rate of eDNA per gram of body mass) tends to decline (Maruyama et al. 2014; Takeuchi et al. 2019; Stoeckle et al. 2021; Yates et al. 2021a, 2021c). The primary mechanisms driving this relationship are likely related to long-recognized allometric scaling in the relationship between body mass and surface area (Meeh 1879; O’Shea et al. 2006), as well as allometric scaling in key physiological rates (*e*.*g*. consumption, excretion, egestion) related to eDNA production (Post et al. 1999; Allegier et al. 2015; Vanni and McIntyre 2016; Yates et al. 2021a). Most of the previous studies examining allometry in eDNA production, however, have focused on *intra*-specific allometry in eDNA production, quantifying how eDNA production changes as body mass increases *within* a species using single-species assays that estimate the concentration of eDNA in an environment using quantitative PCR (qPCR) or digital droplet PCR (ddPCR) methods However, metabolic theory postulates that allometry in key physiological rates also operates at an *inter*-specific scale; organisms from small-bodied species have, on average, higher physiological rates (Brown et al. 2004; Allegier et al. 2015; Vanni and McIntyre 2016) and relationships between body mass and surface area also follow similar patterns across species (Meeh 1879; Reynolds 1996). Yates et al. (2021a) therefore also postulated that eDNA production is likely to follow a similar pattern on an *inter*-specific basis. The extent to which the distribution of eDNA varies interspecifically with organism abundance remains relatively understudied.

Metabarcoding approaches, when applied to eDNA samples, can be used to quantify community species composition (Taberlet et al. 2012). Metabarcoding uses conserved primers to amplify loci across taxonomic groups; amplicons are then sequenced on high-throughput sequencing (HTS) platforms to produce millions of reads that are compared to a reference database for taxonomic identification (Taberlet et al. 2012; Cristescu 2014). A number of studies have demonstrated positive correlations between the relative abundance of species in an environment and relative sequence read counts obtained from metabarcoding (Evans et al. 2016; Hanfling et al. 2016; Lamb et al. 2019; Lawson Handley et al. 2019; Rourke et al. 2021). Although relationships between metabarcoding sequence count and organism abundance are likely to be weaker than for single-species approaches that directly quantify template eDNA concentrations (Yates et al. 2021b) consistent positive correlations between metabarcoding data and relative species abundance have been consistently demonstrated, highlighting its potential utility to infer abundance even if only qualitative or relative abundance comparisons are possible (Kelly et al. 2019; Lamb et al. 2019; Rourke et al. 2021; Yates et al. 2021b).

Relationships between eDNA metabarcoding data and organism abundance could potentially be further strengthened by applying our growing understanding of the ‘ecology of eDNA’ in natural ecosystems. By integrating eDNA dynamics into modelling efforts, researchers may be able to account for some of the unexplained variation in observed relationships between eDNA signal strength estimated by metabarcoding and organism abundance in nature (Yates et al. 2021b). A better understanding of processes involved in eDNA production, for example, could help improve our understanding of the distribution of eDNA observed among species in natural ecosystems.

Metabarcoding datasets derived from sampling natural ecosystems that are paired with simultaneous observational estimates of organism abundance represent a valuable opportunity to empirically estimate how eDNA production scales across species in natural ecosystems. Here we use paired traditional trawling and eDNA metabarcoding data previously published by Stoeckle et al. (2021) to empirically estimate the rate at which eDNA production changes with body mass across species. We applied both frequentist and Bayesian modelling techniques to empirically estimate the inter-specific allometric scaling coefficient (and quantify uncertainty around it) for eDNA production for two groups of fishes represented in the dataset: Osteichthyans (bony fishes) and Chondrichthyans.

## Materials and Methods

### Trawl survey and eDNA collection, extraction, and analysis

For a full description of methodologies used to collect the data, please refer to the original manuscript. In brief, eDNA sampling was paired with trawl data collected off the northeastern coast of the United States for 1-week periods in January, June, August, and November in 2019. A minimum of ten ‘tows’ per three depth intervals for each sampling period were conducted, with a total of 30 traditional trawl samples collected in January and 39 in the other three months. For each tow, species identification, number of individuals, and total weight per species data were collected. Species-accumulation-curves were used to assess whether trawling captured most available species; SACs were calculated for each month using the R package *vegan* (Oksanen et al. 2020). Visual saturation of SACs indicated that trawling likely captured most available species (supplementary files, section 1.0).

For each monthly sampling period, two 1L water samples (one surface and one bottom-depth sample) were collected immediately prior to trawl samples at ten selected tow sites, for a total target of 20 eDNA samples per sampling month period. However, due to bottle breakage and equipment failure, an average of 17 water samples were collected each monthly sampling period. Water samples were maintained on ice and then placed in a -20 °C freezer within 24 h of collection, after which they were thawed and filtered using a 0.45 µm nitrocellulose filter, which was stored in a -80 °C freezer. DNA was subsequently extracted from the filters using a modified PowerSoil kit (Qiagen) (as described in Stoeckle et al. 2020) and extracts were kept frozen prior to library preparation.

DNA processing and bioinformatics were conducted as described in Stoeckle et al. (2020) and Stoeckle et al. (2021). Separate primer sets that target a ∼106-bp segment on the mitochondrial 12S V5 region were used to amplify bony fishes (*Osteichthyes*) and Cartilaginous fishes (*Chondrichthyes*); each eDNA sample was amplified twice: once with bony fish primer set and once with Cartilaginous fish primer set. Bony and Cartilaginous fish amplifications were indexed separately. Sequencing was performed on an Illumina MiSeq for a total of 136 samples (*Chondrichthyan* and bony fishes per field sample) and 79 negative controls. Bioinformatics analysis was conducted using *DADA2* (Callahan et al. 2016), with taxonomic assignments based on a 100% amplicon sequence variant (ASV) match to a 12S reference library of regional fishes (Stoeckle et al. 2017).

### Trawl and eDNA data curation

As in Stoeckle et al. (2021), monthly catch weights were normalized by converting catch data to per-tow values; this facilitated comparisons across sampling periods with different total numbers of tows. Similarly, read numbers were normalized by converting read counts to per-sample values and multiplying by twenty in order to facilitate comparisons across sampling periods in which fewer than 20 samples were collected and analysed. All comparisons were done using monthly trawl weights (sum of normalized trawl catches) and monthly eDNA reads (sum of normalized read counts).

Data for Cartilaginous and bony fishes were analyzed separately for both biological and molecular analysis reasons. First, these two groups have fundamentally different morphological and physiological characteristics that could affect relative eDNA production rates. The surface of *Chondrichthyans*, for example, is characterized by distinct scale morphology and thinner mucous layers compared to bony fishes (Ankhelyi et al. 2018), they possess a distinct excretory system (retention of urea, rectal gland, etc.) (Evans 2010), and they possess morphologically distinct digestive systems (*e*.*g*. spiral valve intestines) (Wetherbee and Gruber 1993). Furthermore, *Chondrichthyan* DNA was amplified using a different primer set; variation in amplification efficiency between the different primer sets could also impact relative recovery of metabarcoding reads. Collectively, differences in relative eDNA production rates and metabarcoding read recovery could significantly affect the estimation of allometric scaling in eDNA production if data from both taxonomic groups are pooled, particularly given that Chondrichthyan species represented in this dataset were, on average, substantially larger-bodied than bony fishes (mean body size = 11.72 vs. 0.63 kg, respectively). Sea lampreys (*Petromyzon marinus*) were excluded due to their ancient divergence from both taxonomic groups (Gess et al. 2006).

For our analyses, we only considered data for fish species that were detected by eDNA sampling in at least one month of the year (i.e. species that were only ever caught in trawls were excluded). Among bony fishes, Northern searobin (*Prionotus carolinus*) represented a significant outlier datapoint, with relative biomass catch in the month of August seven-times higher than the next highest-catch bony fish species across all sampling periods (111 kg-per-tow vs. 16 kg-per-tow, respectively). Preliminary analyses demonstrated that this datum was significantly driving observed relationships and allometric scaling coefficient estimates. The relationship between metabarcoding and original eDNA template concentration can potentially exhibit high residual error due to differences in amplification efficiency across taxa (Elbrecht and Leese 2015; Piñol et al. 2015; Kelly et al. 2019). Similarly, catch per unit effort (CPUE) can also be a poor index of organism abundance under some circumstances (*e*.*g*. hyperstability or hyperdepletion) (Rose and Kulka 1999; Harley et al. 2001; Hubert et al. 2012). Across a large number of datapoints, correlations between these indexes (metabarcoding reads and CPUE and biomass per unit effort (BPUE)) and the underlying variables they track (eDNA template concentration and organism abundance) are likely to be positive in many scenarios. However, a single outlier datapoint exhibiting a disproportionate effect on the analysis could result from stochastic measurement error associated with index variables in relation to the corresponding indicator variables. For the analyses presented in the main text, Northern searobin were therefore excluded; results with this species retained are presented in supplementary materials for comparison.

### Statistical Analyses

#### Integrating allometry into metrics of abundance

Environmental DNA production was assumed to scale allometrically according to the following formula:

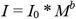

Where *I* = eDNA production rate, *M* = the individual body mass of an organism, I_0_ = a normalization constant, and *b* = an allometric scaling coefficient.

Allometry was integrated into metrics of fish abundance using the following formula:

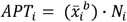

Where *APT*_*i*_ = allometrically scaled abundance-per-tow for the *i*th species, 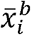 = the mean individual body mass of the *i*th species, *N*_*i*_ *=* the mean capture-per-tow of the *i*th species, and *b* = the interspecific allometric scaling coefficient. Note that a value of 0 for *b* corresponds exactly with species-counts-per-tow, and a value of 1 for *b* corresponds exactly with species biomass-per-tow.

We lacked individual size data for every individual fish, with only total biomass per species and species counts available from tow data which required to use the simplifying assumption that each individual organism could be represented by the mean mass for that taxon (*i*.*e*., Biomass-per-tow (BPT) divided by Individuals-per-tow (IPT)). Although this simplifying assumption may not be suitable across all ecosystems, it is likely a reasonable approximation for our study system. First, in our dataset interspecific variation in body mass is much greater than intra-specific variability in body mass (minimum mean mass = ∼1 g (Bay anchovy, *Anchoa mitchilli*), maximum mean mass = 44 kg (Atlantic sturgeon, *Acipenser oxyrhynchus*). Further, simulations indicate that the level of bias introduced by this assumption is likely small (supplementary files, section 2.0). Under a uniform distribution with body mass variation ranging from 1 to 100 (largest individuals are 100× larger than the smallest) we found that mean percent bias was <5%. This simulated uniform distribution has much higher variation than our observed data. We were also able to generate some individual body mass distribution data by examining tows where only one individual was captured and for all 15 species for which individual data was available for >10 individuals across sampling months interspecific variation was less than in our simulated dataset (supplementary materials, section 2.0). Incorporating individual weights may be more important in study systems where interspecific diversity in body mass is low or where there are small numbers of species in which there is considerable overlap in cohorts within habitats (e.g. small freshwater streams).

#### Frequentist model

To estimate the optimal value of the scaling coefficient using frequentist regression approaches, we repeated methodologies used in (Yates et al. 2021a, 2021c). For each taxonomic group (bony fishes and Chondrichthyans) we built linear models of eDNA metabarcoding read counts as a function of allometrically scaled abundance-per-tow (APT^*b*^) with scaling coefficient values (*b*) ranging from 0.00 to 1.00 by 0.01 intervals. We extracted Akaike Information Criterion (AIC; Akaike 1974) for each model; we predict that the distribution of AIC values corresponding to values of *b* between 0.00 and 1.00 should follow an approximate concave parabolic distribution, with a ‘best-fit’ (*i*.*e*. lowest-AIC) scaling coefficient value occurring at the ‘vertex’ of this approximate parabola (Yates et al. 2021a). AIC and model *r*^2^ values were compared for the ‘optimal’ scaling coefficient model and two traditional metrics of organism abundance: density (*i*.*e*. individuals-per-tow, or ‘IPT’) and biomass (*i*.*e*. biomass-per-tow, or ‘BPT’). All analyses were conducted in *R* (Core R Team 2019).

#### Bayesian model

For Bayesian analysis, we considered the regression model below where *c*_O_ is the regression intercept, APT follows the form in Eq. 2, *c*_1_ is the regression slope, and ε is the variance:

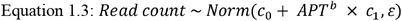

We built separate models for bony fishes and Cartilaginous fishes and implemented Markov Chain Monte Carlo (MCMC) simulations in *JAGS* (Plummer 2003) using the *rjags* (Denwood 2016) and *jagsUI* (Kellner 2021) packages in *R*. For each model, we ran three parallel chains for 500,000 total iterations, discarding the first 5,000 for burn-in and thinning to every 5^th^ iteration to estimate the posterior distribution for each parameter and derive 95% Bayesian Credible Intervals (BCI). We assessed convergence based on a value of R < 1.1 (Gelman and Rubin 1992). We used uniform priors for the latent *b* (0 – 1 for bony fishes and 0 – 3 for Chondrichthyans based on frequentist modeling results; see Results), intercept (0 – 10^6^), beta (0 – 10^5^), and ε (0 – 10^3^ terms) parameters and initial values estimated from the frequentist regression models.

## Results

### Osteichthyes

Bony fish species’ IPT and BPT were significantly and positively correlated with eDNA metabarcoding read counts (Table 2, Figure 1). The distribution of AIC values from the frequentist models exhibited the predicted approximate upward parabolic distribution (Figure 2). The ‘optimal’ AIC value corresponded to a scaling coefficient point-estimate (*b*) of 0.68 with an *r*^2^ = 0.45 and represented a significant improvement over IPT and BPT (ΔAIC = 72.25 and 32.00, *r*^2^ = 0.14 and 0.33, respectively). Similarly, APT^0.68^ exhibited lower Root Mean Square Error (Table 1). Values for *b* between 0.62 and 0.74 had AIC values within two of the lowest AIC value.

**Table 1:** Frequentist results for regressions between eDNA concentration and individuals-per-tow (IPT), biomass-per-tow (BPT), and allometrically scaled abundance per-tow (APT, *b* = 0.68) for bony fish. ΔAIC represents the difference in AIC value between APT^0.68 and the IPT and BPT models. Note that these results are for analyses in which Northern Searobin was excluded.

| Model | $F$ | $P$ | $r^2$ | AIC | $\Delta AIC$ | RMSE |
| --- | --- | --- | --- | --- | --- | --- |
| IPT | 27.45<sub>(1,158)</sub> | <0.001 | 0.14 | 4195.92 | 72.25 | 117541.8 |
| BPT | 80.06<sub>(1,158)</sub> | <0.001 | 0.33 | 4155.96 | 32.00 | 103744.8 |
| APT<sup>0.68</sup> | 133.3<sub>(1,158)</sub> | <0.001 | 0.45 | 4123.67 | - | 93787.5 |

**Table 2:** Frequentist results for regressions between eDNA concentration and individuals-per-tow (IPT), biomass-per-tow (BPT), and allometrically scaled abundance per-tow (APT, *b* = 1.62) for Chondrichthyans. ΔAIC represents the difference in AIC value between APT^0.68 and the IPT and BPT models.

| Model | $F$ | $P$ | $r^2$ | AIC | $\Delta AIC$ | RMSE |
| --- | --- | --- | --- | --- | --- | --- |
| IPT | 8.47<sub>(1,35)</sub> | 0.006 | 0.17 | 1061.00 | 24.53 | 376185.1 |
| BPT | 21.22<sub>(1,35)</sub> | <0.001 | 0.36 | 1051.48 | 15.01 | 330766.4 |
| APT<sup>1.62</sup> | 49.37<sub>(1,35)</sub> | <0.001 | 0.57 | 1036.47 | - | 270024.2 |

**Figure 1:**
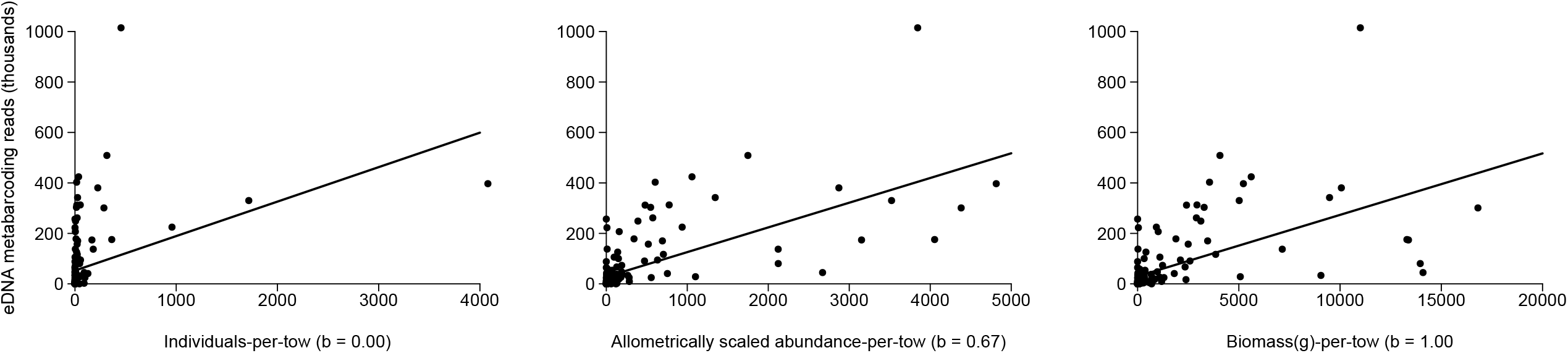
Linear regressions for bony fish species’ eDNA metabarcoding read count (thousands) and three metrics of abundance: (a) Individuals-per-tow, (b) Allometrically-scaled abundance-per-tow (*b* = 0.67) (APT^0.67^), and (c) Biomass-per-tow. Note that the scaling coefficient estimate represented in figure (b) was the point-estimate derived from the Bayesian model.

**Figure 2:**
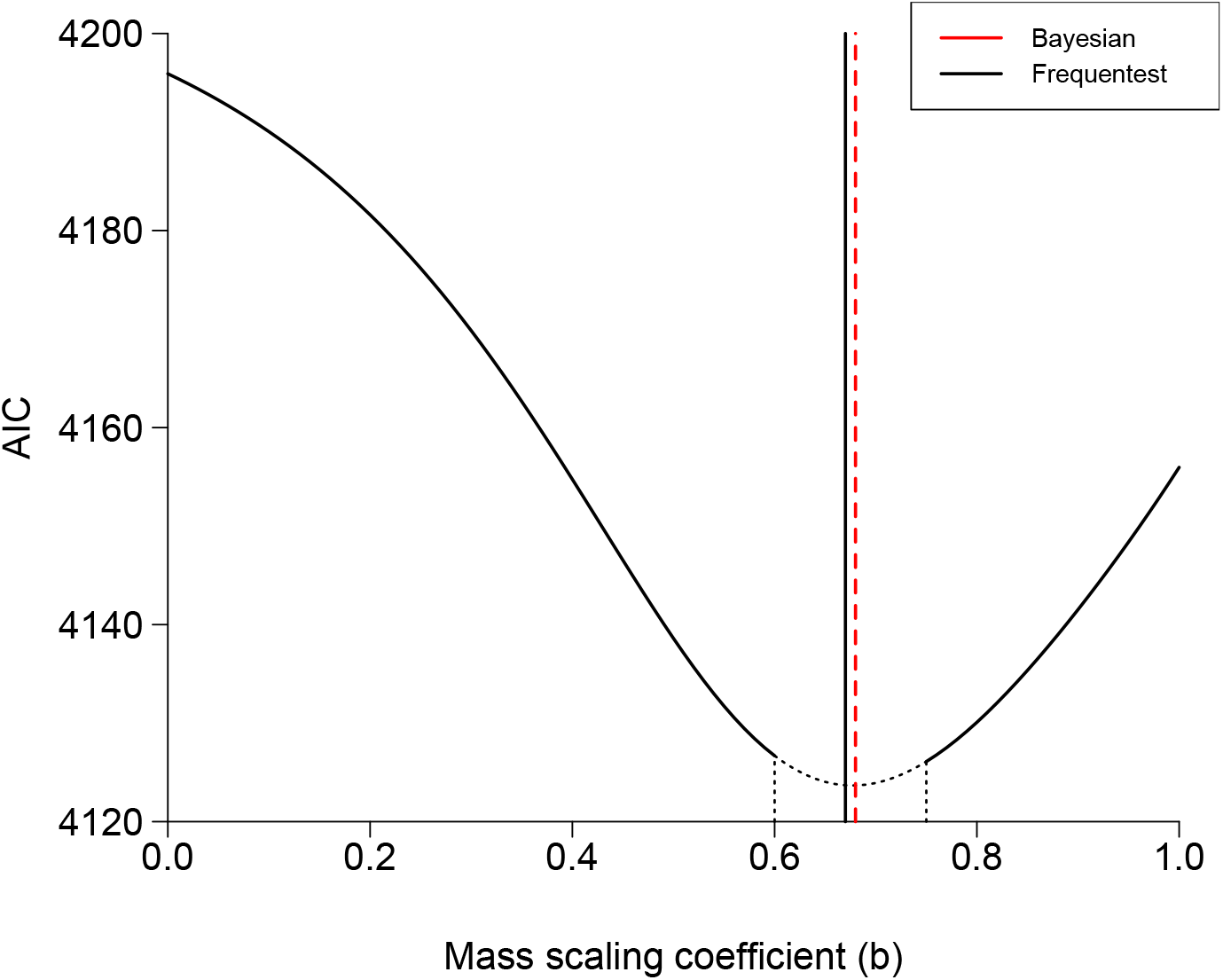
Distribution of AIC values for linear regressions between bony fish abundance and eDNA metabarcoding read count, corresponding to scaling coefficients ranging from 0.00 to 1.00. The dotted portion of the curve denoted with vertical dotted lines denotes the range of models with ΔAIC < 2 of the ‘optimal’ scaling coefficient model. The black vertical line represents the best-fit scaling coefficient estimated using frequentist approaches, and the red vertical line represents the best-fit scaling coefficient estimated using the Bayesian model.

Values from the frequentist model with a scaling coefficient of 0.68 were used to initialize parameter values for the Bayesian model, although model results were largely unaffected by initial parameter values. The Bayesian model converged successfully and we derived a point estimate for the scaling coefficient of 0.67 with a 95% Bayesian credible interval of 0.59 – 0.77 (Figures S1), exhibiting close correspondence to the frequentist approach (Figure 2).

Analyses above removed Northern Searobin, which was a high leverage, outlier datum. Analyses of bony fish data with Northern Searobin included still found evidence of allometric scaling; the point estimates for the scaling coefficient using frequentist approaches and Bayesian methods were both 0.55 (credible interval = 0.46 – 0.67) (see supplementary materials, section 3.0). Allometric scaling in eDNA production was therefore still observed/inferred regardless of whether this outlier species was included, but the point estimate and credible interval range for *b* were substantially lower.

### Chondrichthyes

Chondrichthyan species’ IPT and BPT were significantly and positively correlated with eDNA metabarcoding read counts (see Table 2, Figure 3). The distribution of AIC values from the frequentist models with scaling coefficient values ranging from 0.00 to 1.00 did not exhibit the predicted upward parabolic distribution; AIC values instead declined as scaling coefficients approached BPT (*b* = 1.00). We therefore explored the effect of scaling coefficient values > 1.00. The predicted ‘approximate’ upward parabolic distribution of AIC values was only observed when scaling coefficient values were extended to 2.00 (Figure 4). The ‘optimal’ AIC value observed corresponded to a scaling coefficient value of 1.62 with an *r*^2^ = 0.57, implying that large-bodied *Chondrichthyans* had higher mass-specific eDNA production rates than smaller *Chondrichthyans*; this model represented a significant improvement over IPT and BPT (ΔAIC = 19.25 and 11.69, *r*^2^ = 0.19 and 0.36, respectively).

**Figure 3:**
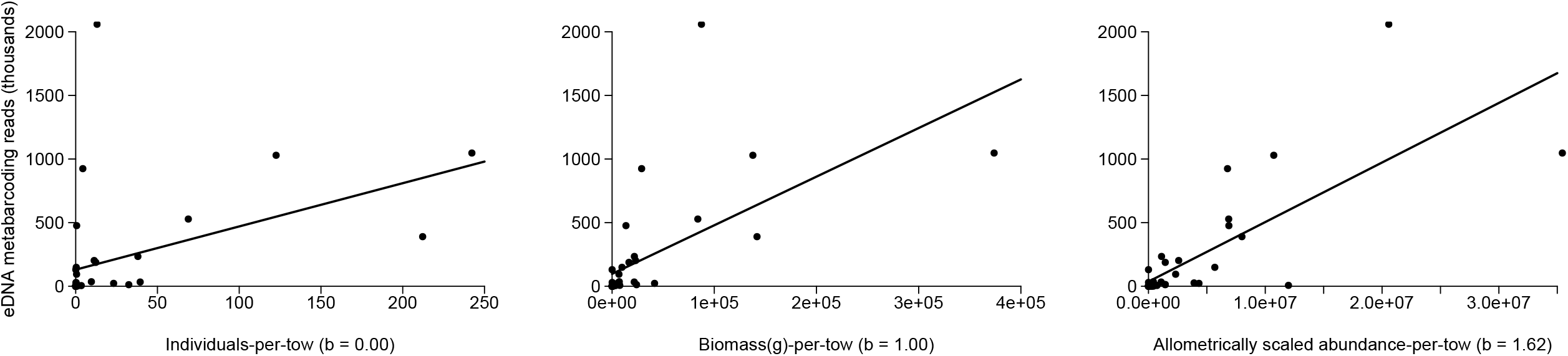
Linear regressions for Chondrichthyan species’ eDNA metabarcoding read count (thousands) and three metrics of abundance: (a) Individuals-per-tow, (b) Allometrically-scaled abundance-per-tow (*b* = 1.62) (APT^1.62^), and (c) Biomass-per-tow. Note that the scaling coefficient estimate represented in figure (b) was the point-estimate derived from the frequentist model as our Bayesian model failed to converge.

**Figure 4:**
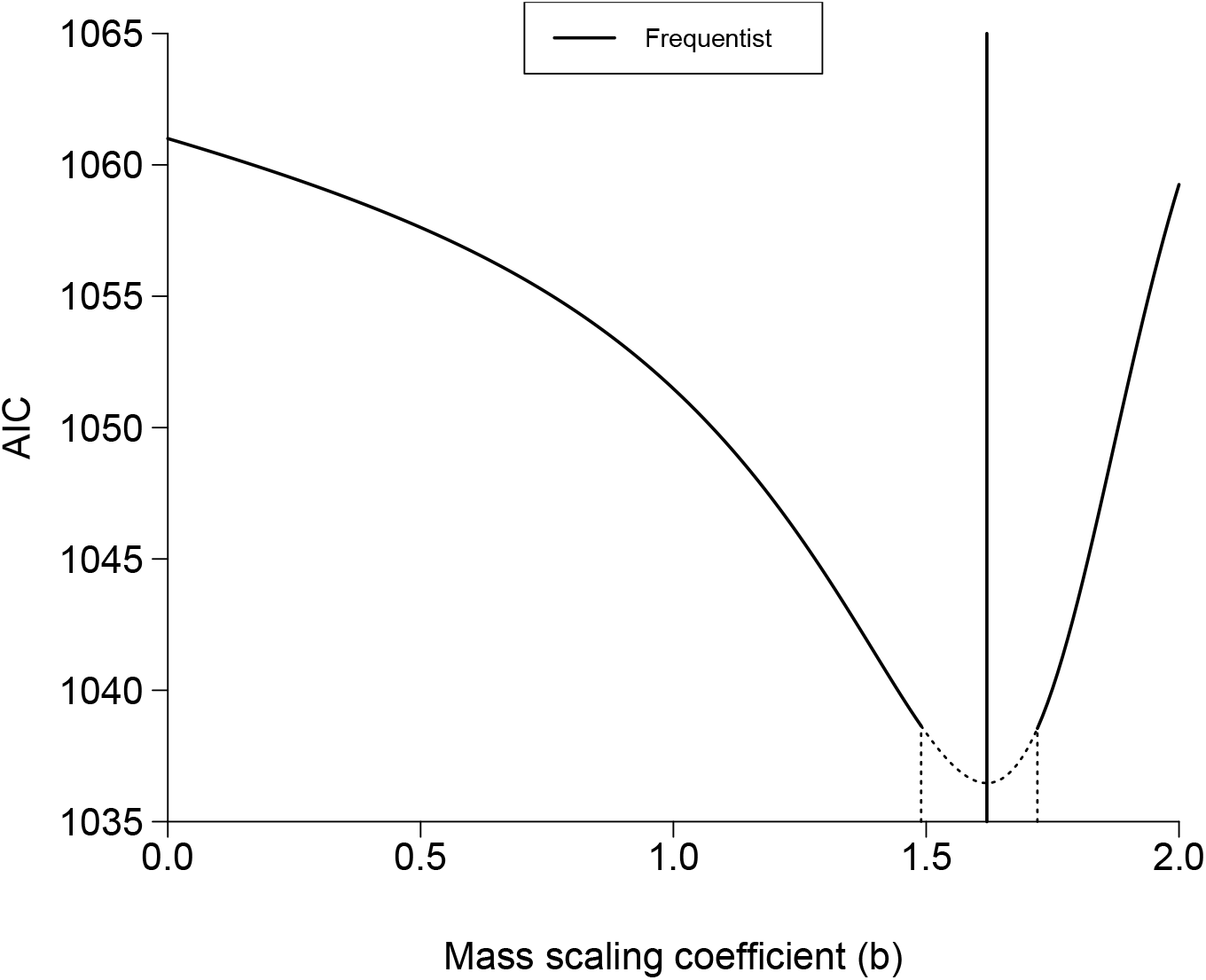
Distribution of AIC values for linear regressions between Cartilaginous fish abundance and eDNA metabarcoding read count, corresponding to scaling coefficients ranging from 0.00 to 2.00. The dotted portion of the curve denoted with vertical dotted lines denotes the range of models with ΔAIC < 2 of the ‘optimal’ scaling coefficient model. The black vertical line represents the best-fit scaling coefficient estimated using frequentist approaches; note that the Bayesian model point estimate was 1.52, but the model failed to converge.

Values from the frequentist model with a scaling coefficient of 1.62 were used to initialize parameter values for the Bayesian model. We derived a point estimate for the scaling coefficient from the Bayesian model of 1.52, a considerable discrepancy from the frequentist model (Figures 4 and S2). The Bayesian model, however, did not converge successfully regardless of initialized parameter values and the posterior distribution of the MCMC estimate for *b*_*chond*_ exhibited a bimodal distribution that was variable between chains (Figure S2).

## Discussion

We found that eDNA shedding rates scale allometrically (i.e., larger fish produce less eDNA per unit body mass) across bony fish species in a natural system. This is an important observation because when researchers correlate quantitative eDNA data with traditional metrics of organism abundance (e.g. numerical abundance or biomass) they make implicit assumptions about the value of the allometric scaling coefficient (*b*). Further research is needed to test generality across systems, but this work e demonstrates 1) that it is possible to infer interspecific allometric scaling coefficients from large observational datasets of paired eDNA metabarcoding and animal capture data in natural ecosystems and 2) that incorporating an understanding of allometric scaling may strengthen correlations between read count data and organism abundance. Although inter-specific allometry in eDNA production should also be experimentally validated under controlled conditions, large-scale observational datasets represent valuable opportunities to explore the fundamental ‘ecology’ of eDNA production in natural ecosystems. Future large-scale studies should similarly attempt to infer whether allometrically scaling processes may be affecting the distribution of eDNA in natural ecosystems. Furthermore, while common Bayesian modelling packages can provide the modelling flexibility to directly estimate non-linear model parameters and their error, they are also useful because ‘informative priors’ can be used to shape models when insufficient data are available to directly estimate coefficients. Smaller-scale studies that lack the sample size to accurately estimate scaling coefficients directly could potentially use informative priors based on previous work and potentially strengthen resulting eDNA/abundance correlations.

We do note, however, that we anticipate that the value of integrating allometry to abundance/eDNA concentration correlations will be lower with smaller metabarcoding datasets (*e*.*g*. < ∼15 species). A number of factors (e.g. differences in PCR efficiency among species (Elbrecht and Leese 2015; Leese et al. 2021), community composition (Piñol et al. 2019)) can potentially introduce substantial ‘noise’ into the relationship between quantitative eDNA read data and organism abundance (Kelly et al. 2019; Yates et al. 2021b). In large datasets (i.e. > ∼15-20 species), such ‘noise’ likely ‘averages out’; in small datasets, stochastic idiosyncrasy in amplification efficiency among taxa may well ‘drown out’ allometric effects of eDNA production. Advances in metabarcoding that improve the relationship between read count and original template eDNA concentrations (e.g. correcting for taxa-specific amplification efficiency) could potentially improve the utility of correcting for allometry (Kelly et al. 2019). At the very least, however, future studies should explore the utility of transforming abundance data using scaling coefficients derived from theoretical expectations; a scaling coefficient value of approximately 2/3 appears to be both theoretically and empirically justified.

We found strong empirical evidence that the value of *b* for bony fish eDNA production corresponds closely to theoretical expectations based on whole-body surface area and physiological allometric scaling. Stoeckle et al. (2021) investigated the effect of allometry on eDNA production by fixing the value of *b* in their ‘allometric abundance index’ to 2/3 based on these theoretical expectations; this appears to have been a well-justified approach, given that our empirically-derived point-estimate for the value of *b* for bony fishes corresponded to 0.67. Our Bayesian model also provided a robust empirical estimate of uncertainty around this variable, with a credible interval value of 0.58 – 0.77, although it is important to note that this is likely a ‘liberal’ estimate of the credible interval due to the fact that the independent variable in the analysis (organism abundance) was also estimated with error, rather than experimentally controlled (e.g. as in a mesocosm experiment). Notably, the point-estimate and credible interval corresponded to allometric scaling rates observed for body surface area in fish (O’Shea et al. 2006) and for key physiological rates like consumption/excretion/egestion (Post et al. 1999; Wiff and Roa-Ureta 2008; Allegier et al. 2015; Vanni and McIntyre 2016). However, metabolic allometry does not appear to be driving the relationship, with upper credible intervals barely overlapping with classic theoretical scaling relationships postulated from metabolic theory (e.g. *b* = 0.75) (Brown et al. 2004) and non-overlapping with a recently derived empirical estimate of 0.89 (0.82 – 0.99) for the value of the metabolic scaling coefficient in fishes (Jerde et al. 2019). These differences were accentuated if data for Northern Searobin (an outlier species) were retained in the analyses.

Understanding the physiological processes involved in eDNA production also has potential implications for the application of eDNA for monitoring. Although model predictive power was low overall (r^2^ = 0.45), integrating allometry (i.e. APT^0.67^) linearized eDNA/abundance data (see Figure S3) and significantly improved model predictive capacity, as indicated by RMSE values, relative to models based on traditional metrics. The linear relationship between organism abundance and read count we observed was weaker than typically observed for single-species qPCR/dPCR approaches (Yates et al. 2019), likely due to (previously discussed) factors in metabarcoding that can introduce ‘noise’ in the relationship between read count and template eDNA concentrations (Yates et al. 2021b). Our results, however, suggest that the physiology of eDNA production remains an important consideration for metabarcoding studies attempting to infer relative organism abundance from eDNA read count.

Scaling factors and constants, however, may need to be specific to taxonomic/phylogenetic groups because physiological processes involved in eDNA production are complex and likely variable across species and taxonomic groups (Sassoubre et al. 2016; Yates et al. 2021b). Bony fishes and Cartilaginous fishes, for example, exhibit substantial differences in physiology and morphology that could impact relative eDNA production and, thus, cross-taxonomic allometric relationships in eDNA production. Similarly, physiological processes among specific taxa could also result in a breakdown of typical allometric relationships in eDNA production. Reproductive activity (e.g. broadcast spawning) can result in an increase in mass-specific eDNA production rate (Takeuchi et al. 2019; Curtis et al. 2020); accounting for reproductive activities during eDNA sample collection timing could be relevant, particularly for species that might appear as ‘outliers’ with larger-than-expected mass-specific eDNA production rates. Variability in primer amplification efficiency (both within and between primer sets) might also limit inter-taxa comparisons between read counts and organism abundance (Elbrecht and Leese 2015; Kelly et al. 2019). Accounting for phylogeny may therefore be important when conducting such comparisons across broad taxonomic groups.

While bony fishes exhibited patterns almost exactly corresponding to theoretical expectations, Chondrichthyans deviated substantially from predictions, with frequentist approaches provided a scaling-coefficient estimate of 1.62 and our Bayesian models failing to converge. While this may be indicative of a potential biological phenomenon warranting further study, the more likely explanation is that we simply lacked the statistical sample size/power to estimate the value of the scaling factor from potentially ‘noisy’ metabarcoding/trawl data; with only 13 species and 37 datapoints, the Chondrichthyan dataset had significantly lower representation relative to bony fish (with 56 species with 160 datapoints). Variability among amplification efficiency can be common even among closely related taxa in metabarcoding (Elbrecht and Leese 2015; Piñol et al. 2015), introducing potential residual error in the relationship between metabarcoding read count and organism abundance due to imperfect correlation between final read count and the original template eDNA concentrations. Catch-per-unit-effort (CPUE) data can also be poorly correlated with abundance for some species and systems (Rose and Kulka 1999; Harley et al. 2001; Hubert et al. 2012; Yates et al. 2021a). As a result, error between our index variables (read count and CPUE/BPUE data) and underlying fundamental parameters (eDNA concentration and organism abundance), combined with small sample size, could potentially account for both the unusual results we observed for Chondrichthyans and the failure of our Bayesian model to converge even with model parameters initialized based on the frequentist model.

## Conclusions and Recommendations

Our findings demonstrate that considering the physiology of eDNA production may be important for the future application of eDNA sampling to monitor organism abundance. Understanding the consistency of the effect of allometric processes on eDNA production, as well as conditions under which allometric patterns might emerge, is crucial for evaluating the extent to which integrating allometry can improve eDNA/abundance correlations and, ultimately, help operationalize eDNA to monitor abundance and biodiversity in natural ecosystems. The extent to which our findings might apply to other ecosystems remains unknown. We encourage future studies to consider the implicit assumptions regarding the physiology of eDNA production made correlating eDNA with species *N* and biomass.

We also want to re-emphasize that correlating quantitative eDNA data exclusively to traditional metrics of organism abundance (*e*.*g. N* and biomass) makes inherent assumptions regarding the physiology of eDNA production, regardless of whether researchers explicitly consider or evaluate the value of the allometric scaling coefficient (*b*) within their own study systems; comparing eDNA to *N* or biomass simply fixes the value of *b* at 0 or 1, respectively. Future researchers should explicitly consider and evaluate how they might expect eDNA production to scale allometrically within and across species – is eDNA production likely to be a function of numerical abundance *(i*.*e*., *b* = 0), biomass *(i*.*e*., *b* = 1), or something in between (*i*.*e*., 0 < *b* > 1)? In particular, we would encourage future studies to consider allometry in study systems where populations or species exhibit substantial variation in body size distributions; allometry is unlikely to affect relative eDNA production rates when organisms exhibit similar body sizes across taxa. Understanding the consistency of the effect of allometric processes on eDNA production, as well as conditions under which allometric patterns might emerge, is crucial for evaluating the extent to which integrating allometry can improve eDNA/abundance correlations and, ultimately, help operationalize eDNA to monitor abundance in natural ecosystems.

## Supporting information

Supplementary Analyses

## Acknowledgements

We would like to thank Adam Sepulveda, whose review of our manuscript greatly improved subsequent versions and whose ideas and commentary opened up a number of new avenues to explore. MCY was funded by a CIGLR post-doctoral fellowship.

## Supplementary Figures

**Figure S1:**
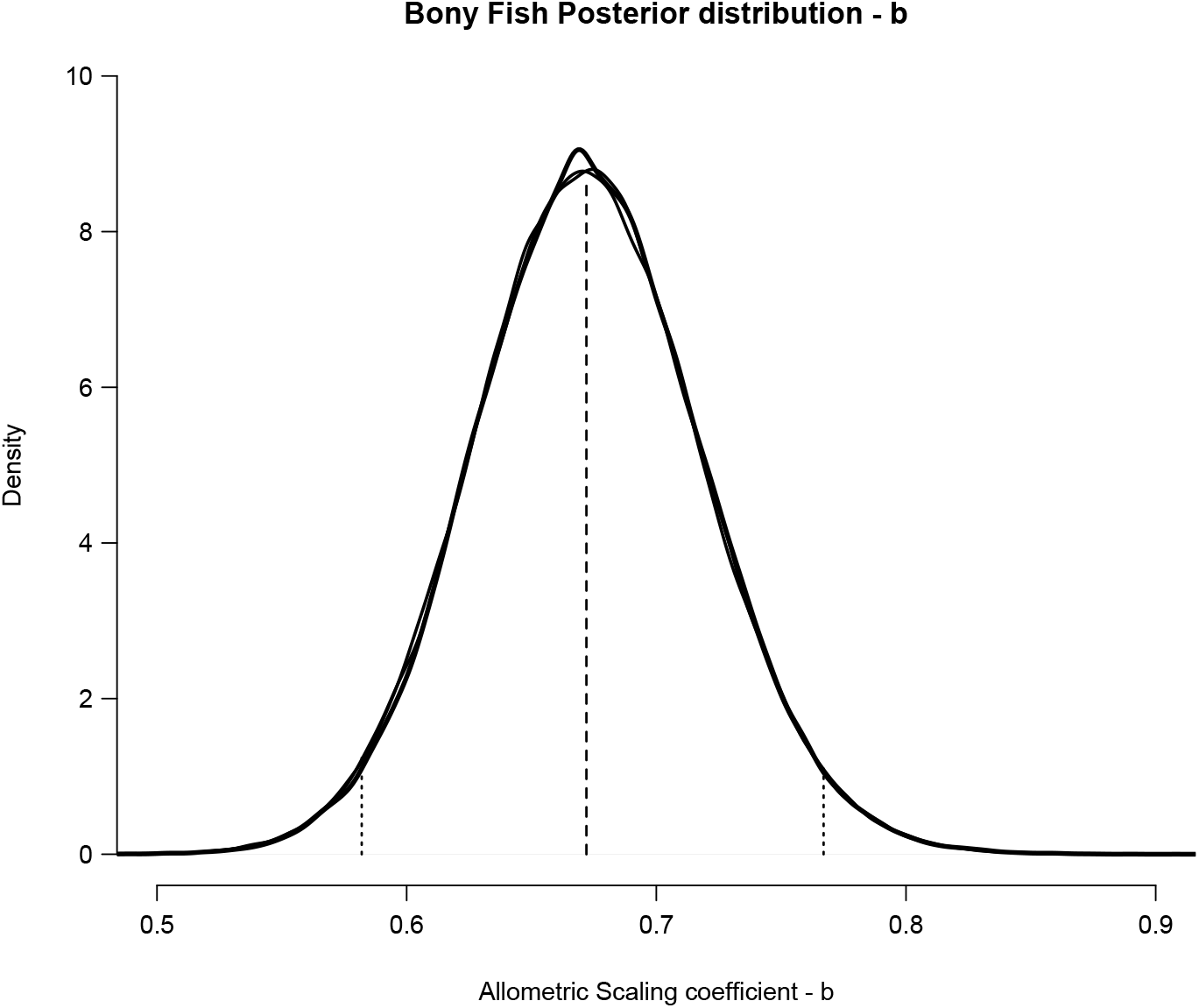
Posterior distribution of the allometric scaling parameter for bony fishes, including median and 95% credible interval.

**Figure S2:**
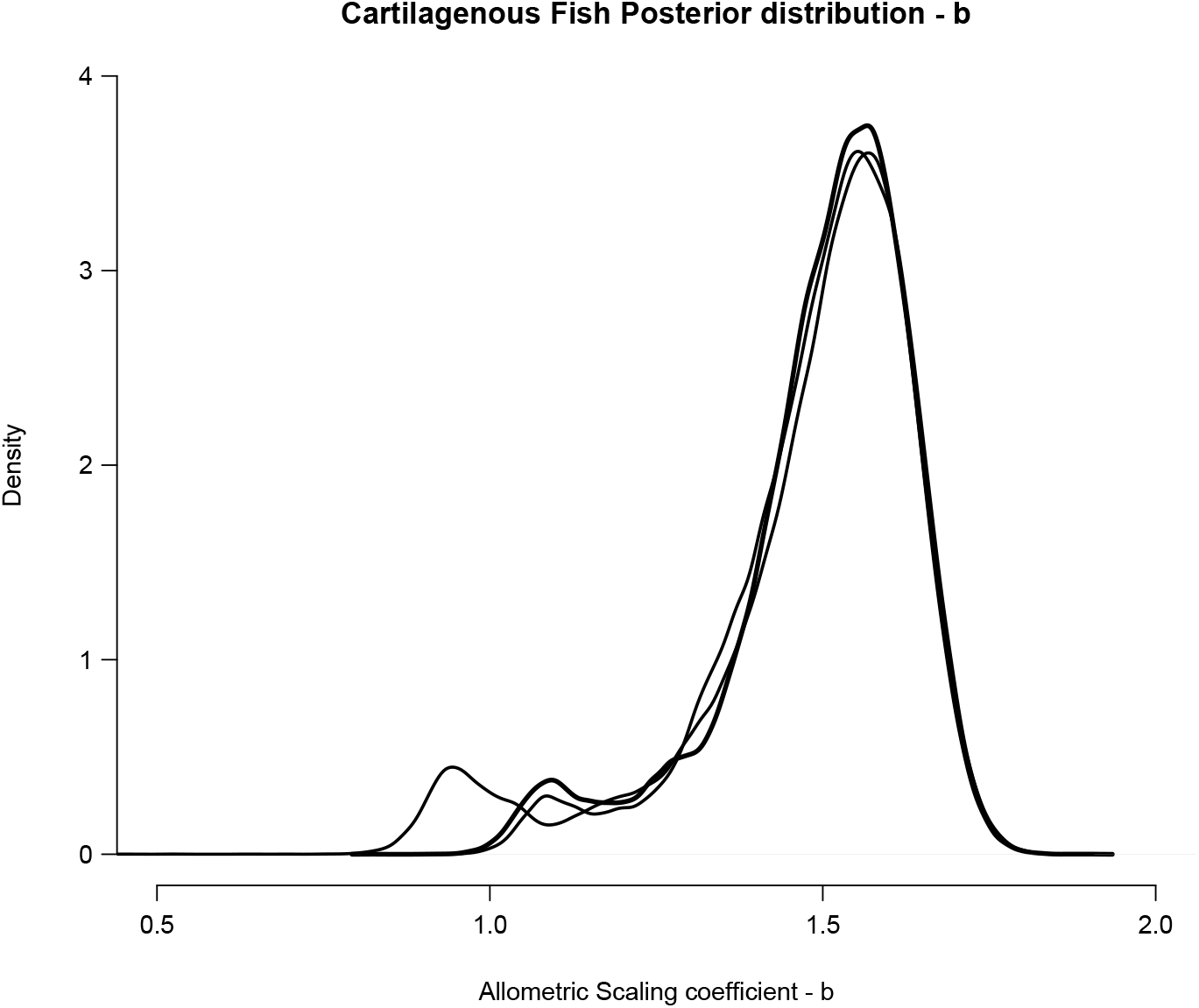
Posterior distribution of the allometric scaling parameter for Cartilaginous fishes; note that model failed to converge.

**Figure S3:**
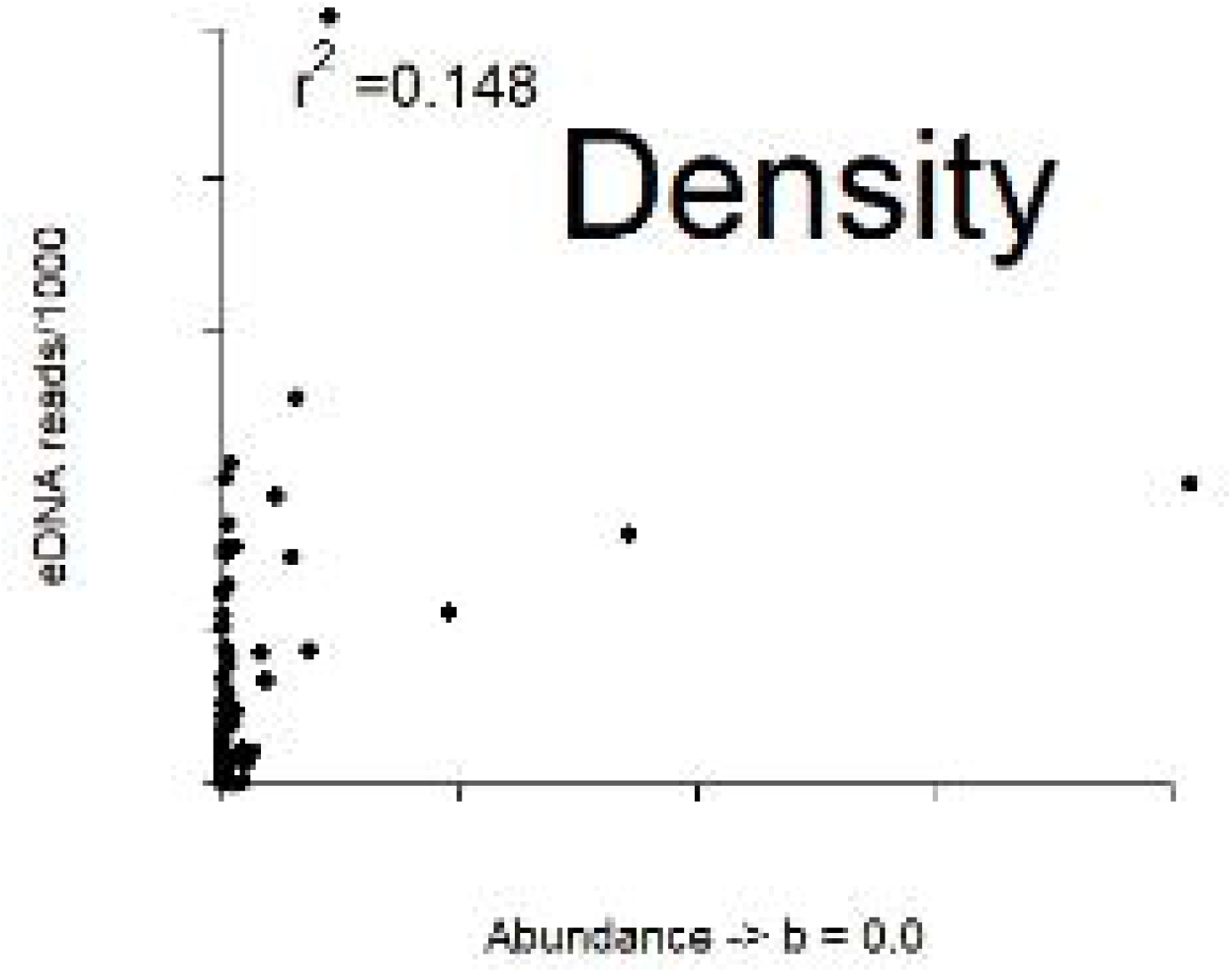
Demonstrating how a gradient of scaling factor values (from ‘b’ = 0 to ‘b’ = 1.0, by 0.1 intervals) affects the linear relationship between organism abundance and metabarcoding read count.

## Notes

### Competing Interest Statement

The authors have declared no competing interest.

