## Supplementary Analyses for "Interspecific allometric scaling in eDNA production in fishes reflects physiological and surface area allometry"

### Appendix

#### Section 1.0: Species accumulation curves for trawl data

We calculated species accumulation curves using 100 permutations of the individual trawl data for each month using the R package *vegan* (Oksanen et al. 2020). Visual saturation of SACs indicate that trawling likely captured most available fish species.

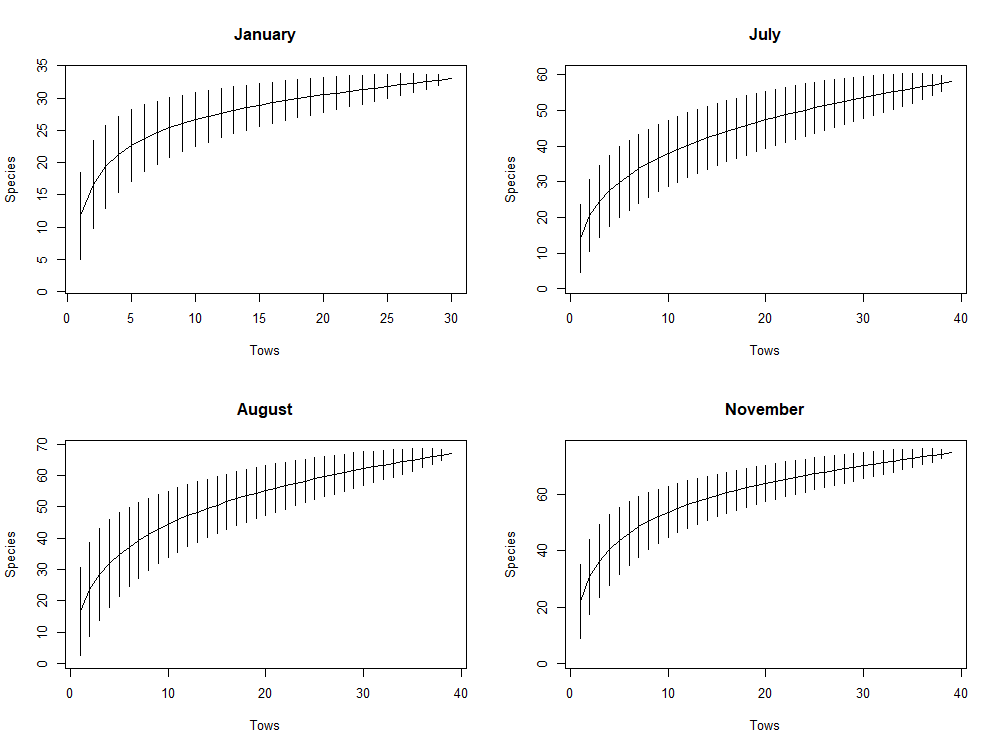

Figure A1: Species accumulation curves for trawl data from each survey month. Error bars represent two standard deviations above and below the point estimate.

#### Section 2.0: Elasticity analysis

**Simulation**

To assess potential bias introduced by failing to account for variation in individual mass, we conducted a simulation. Ten individuals were drawn from a uniform distribution of masses from 1 – 100 (i.e., the maximum possible individual mass is 100× the minimum possible individual mass) and calculated allometrically-scaled mass using a *b* value of 0.67 first accounting for individual mass and then with the simplifying assumption that all individuals were equal to the mean mass. We calculated percent bias (PBIAS) as the difference between the two estimates divided by the estimate that accounts for individual mass (truth). We ran this simulation in 1,000 iterations. Mean percent bias was 4.52% (95% CI 1.12 – 9.59%). The level of individual variation simulated here is substantially higher than that present in our data (see below).

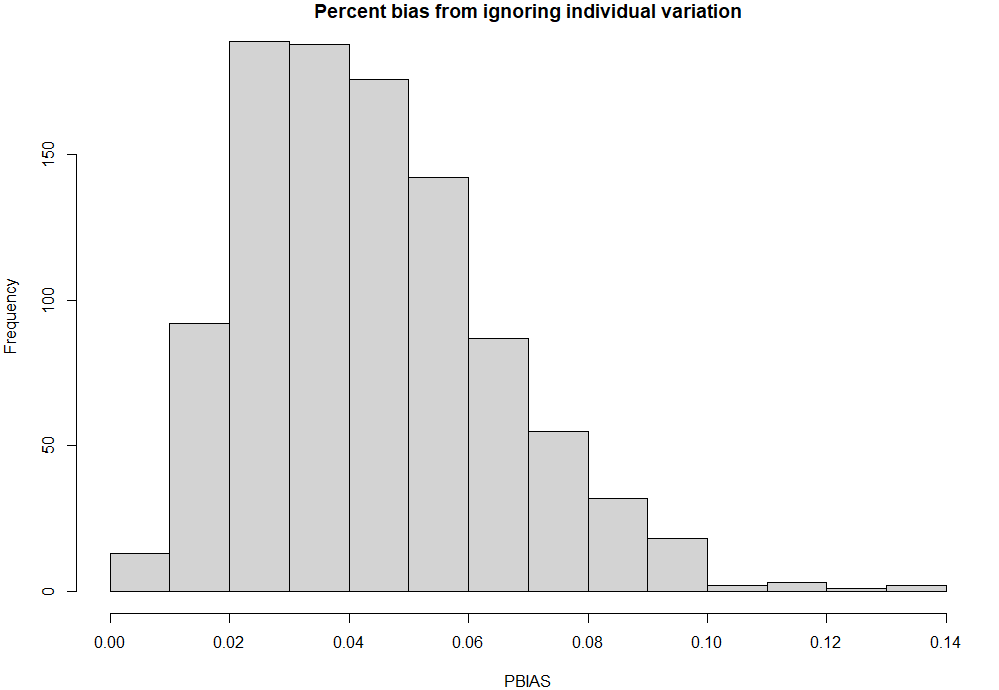

Figure A2: mean percent bias generated by ignoring individual variation when estimating allometrically scaled mass from 10 individuals randomly drawn from a uniform ‘body mass’ distribution ranging from 1-100.

**Empirical data**

Individual weight data was only available when only a single individual of a taxon was captured in a tow (n = 545 individual weights). For 15 taxa, there were individual weights available for >10 individuals across all months.

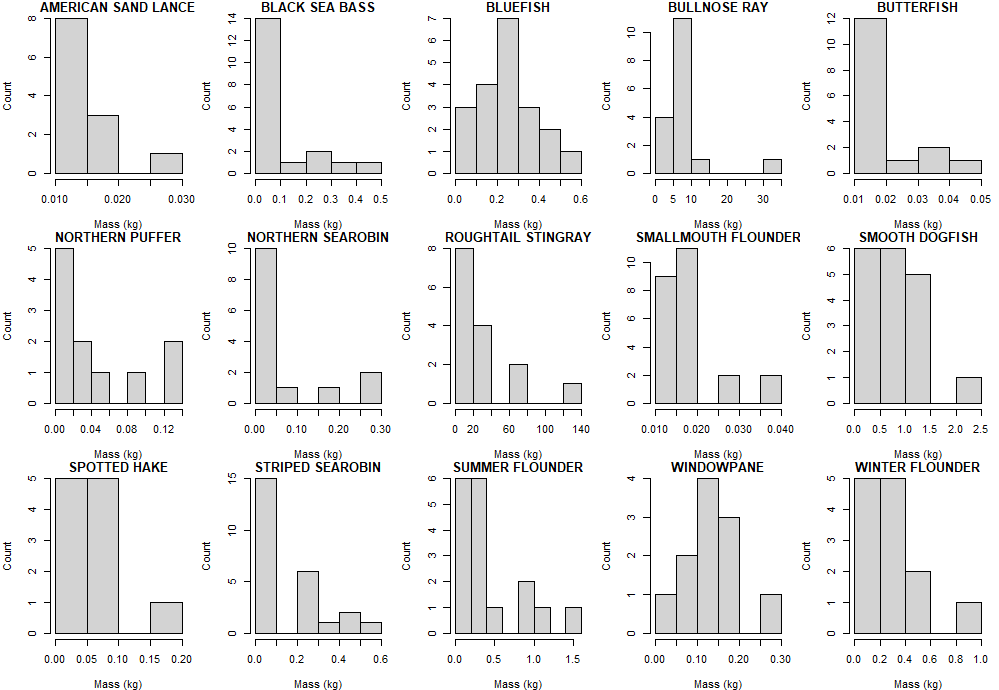

Figure A3: Individual weights for species derived from trawl sets in which only a single individual was captured.

For these 15 taxa we calculated the maximum observed individual mass divided by the minimum observed individual mass. The largest individual ranged from 3.0 – 58.7× that of the smallest (mean = 25.3×). Also note that estimates of mean species’ mass were based on monthly estimates, whereas these values were drawn from all tows across years. As a result, the variation observed above in body mass within a species is likely substantially higher than typically observed within a season, indicating that our allometrically scaled estimates of species biomass would exhibit less bias.

#### Section 3.0: Northern Searobin included in analysis

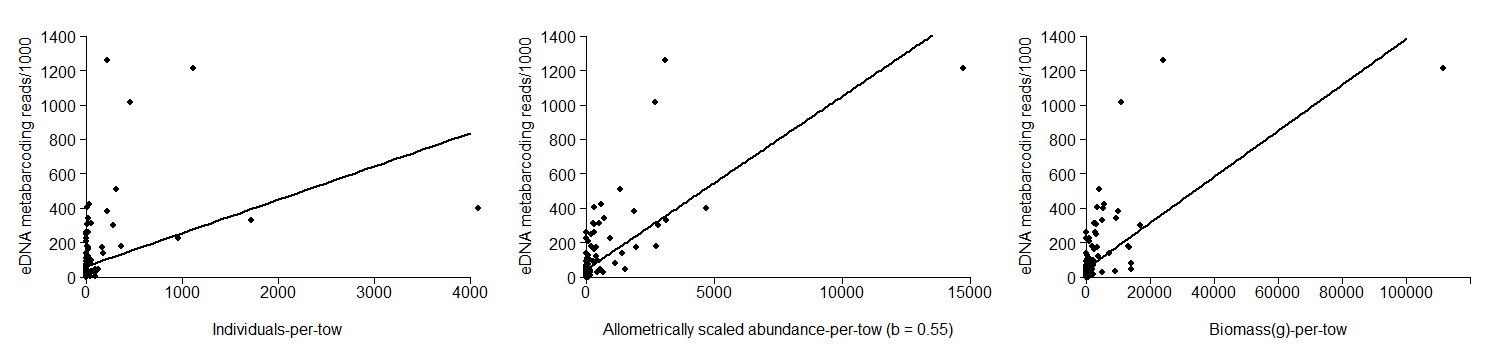
When Northern Searobin are included in the analysis, Bony fish species’ individuals-per-tow (IPT) and biomass-per-tow (BPT) were still significantly and positively correlated with read counts from eDNA (see Table S1, Figure A1). The distribution of AIC values from the frequentist models with scaling coefficient values ranging from 0.00 to 1.00 still exhibited the predicted approximate upward parabolic distribution (Figure A2). The ‘optimal’ AIC value corresponded to a scaling coefficient point-estimate (*b*_OST_) of 0.55 with an *r*^2^ = 0.54, and represented a significant improvement over IPT and BPT (∆AIC = 99.75 and 21.84, *r*^2^ = 0.15 and 0.47, respectively). Values for *b*_Ost_ between 0.49 and 0.63 had AIC values within 2 of the ‘optimal’ AIC value.

Figure A4: Linear regressions for bony fish species’ eDNA metabarcoding read count (divided by 1000) and three metrics of abundance: (a) Individuals-per-tow, (b) Allometrically-scaled abundance-per-tow (*b* = 0.67) (APT^0.55^), and (c) Biomass-per-tow. Note that the scaling coefficient estimate represented in figure (b) was the point-estimate derived from the Bayesian model.

Values from the frequentist model with a scaling coefficient of 0.55 were used to initialize parameter values for the Bayesian model, although model results were largely unaffected by initial parameter values. The Bayesian model converged successfully, and the posterior distribution of the estimate for *b_Ost_* exhibited an approximately normal distribution that was consistent across chains. The point estimate for the scaling coefficient from the Bayesian model was 0.55 with a credible interval of 0.46 and 0.67 (Figures A3), exhibiting close
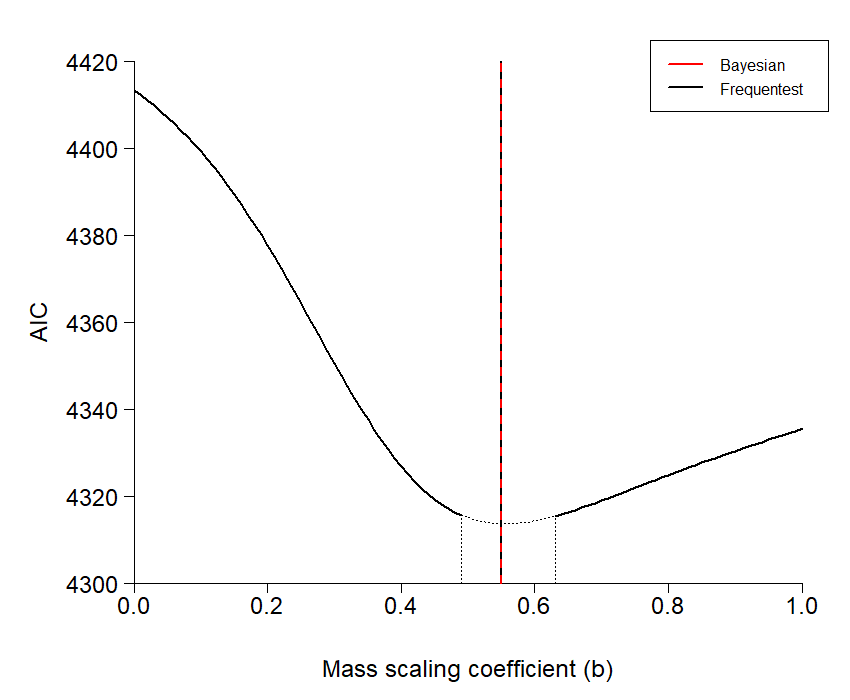
correspondence to our frequentist approach (Figure A2).

Figure A5: Distribution of AIC values for linear regressions between bony fish abundance and eDNA metabarcoding read count with Northern Searobin included, corresponding to scaling coefficients ranging from 0.00 to 1.00. The dotted portion of the curve denoted with vertical dotted lines denotes the range of models with ΔAIC < 2 of the ‘optimal’ scaling coefficient model. The black vertical line represents the best-fit scaling coefficient estimated using frequentist approaches, and the red vertical line represents the best-fit scaling coefficient estimated using the Bayesian model.

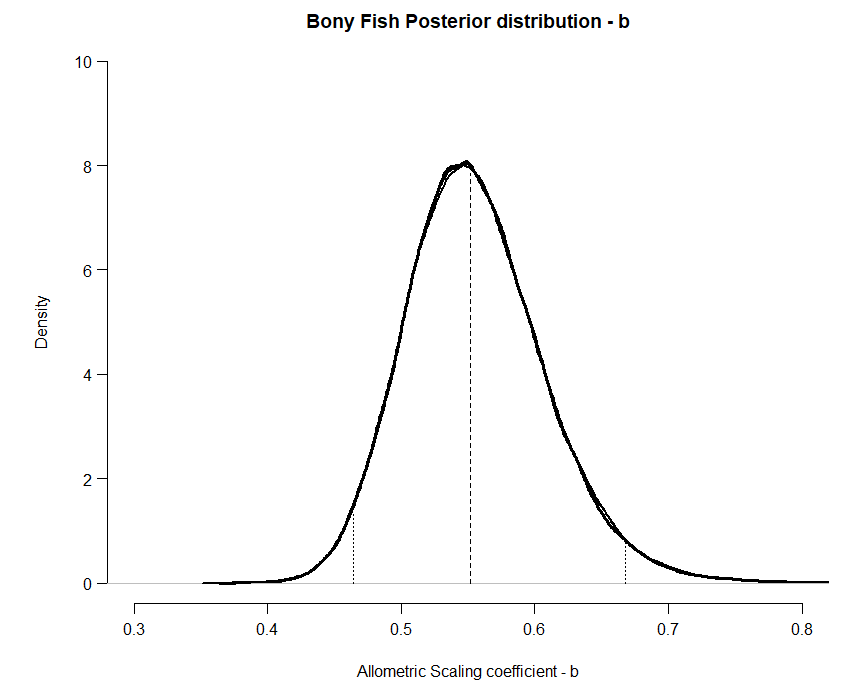
Figure A6: Posterior distribution of the allometric scaling parameter for bony fishes with Northern Searobin included, including median and 95% credible interval.

Table S1: Frequentist results for regressions between eDNA concentration and individuals-per-tow (IPT), biomass-per-tow (BPT), and allometrically scaled abundance per-tow (APT, *b* = 0.55) for bony fish.

| Model | *F* | *P* | *r^2^* | AIC | ΔAIC |
| --- | --- | --- | --- | --- | --- |
| IPT | 29.28_(1,162)_ | <0.001 | 0.15 | 4413.36 | 99.75 |
| BPT | 145.6_(1,162)_ | <0.001 | 0.47 | 4335.45 | 21.84 |
| APT^0.56^ | 189.4_(1,162)_ | <0.001 | 0.54 | 4313.61 | - |
